# KinConfBench: A Curated Benchmark for Cofolding Models on Kinase Conformational States

**DOI:** 10.64898/2026.04.07.716788

**Authors:** Kunyang Sun, Teresa Head-Gordon

## Abstract

Protein kinases are critical drug targets, requiring therapeutics that can modulate their active and inactive conformational states. While cofolding models can generate global folds directly from kinase sequences and ligand SMILES strings, these models have not yet been tested on their ability to recover ligand induced-fit conformational states of the kinase proteins. Here, we introduce KinConfBench, a curated benchmark of 2,225 high-quality human kinase chains to evaluate the ability of three state-of-the-art cofolding models—Boltz-2, Chai-1, and Protenix—to recover both canonical and rare conformational states. We show that geometric success metrics of a ligand pose in the active site does not correlate strongly with the correct kinase conformational state, motivating a new set of dynamical benchmarks for assessing cofolding models. While all three cofolding models achieve ∼65-75% prediction accuracy for kinase conformational classification, they exhibit severe mode collapse when performing multiple inferences, show negligible structural diversity in sampling induced-fit motions, and display a prevalent “apo-drift” in which all three cofolding models predominately predict the kinase to be in its ligand-free state. Our results highlight that capturing ligand-induced protein conformational diversity, not just geometric fit, is critical for next-generation structure-based drug discovery.

## Introduction

Highly accurate data-driven protein structure prediction models [1–6] have become powerful tools for structural biology. By combining evolutionary information with structure-based training using the Protein Data Bank (PDB) [7], these models have allowed researchers to generate plausible protein structural hypotheses from sequence, especially for proteins that are not structurally enabled by experiment. Inspired by this advancement, new directions have opened up to extend the framework beyond single-chain proteins, predicting protein complexes with other proteins[8, 9] or biologics [10–16]. Consequently, a new class of “cofolding” models has emerged, designed to predict structures for biomolecular complexes including proteins, DNAs, RNAs, post-translational modifications, ions, small molecules, and other modalities critical for drug discovery. Models such as NeuralPLexer[10], AlphaFold3[11], Chai-1[13], Protenix[16], and Boltz 1 and 2[14, 15] have shown promise in accelerating structure-based drug discovery (SBDD) campaigns by generating structural hypotheses at a reasonable computational cost.

Despite their rapid adoption, data-driven cofolding models suffer from distinct generalization issues. For instance, Škrinjar et al. observe that these models exhibit significantly lower success rates and confidence scores when tasked with systems outside their training distribution [17]. Beyond these failures in low-data regimes, even predictions with high confidence scores can exhibit physically invalid steric clashes or distorted ligand geometries that fails the benchmark evaluations by the PoseBusters suite [18]. Moreover, recent benchmarking efforts have attempted to probe limitations in other regimes such as covalent modifications [19], PROTACs [20], and molecular glues [21]. Besides these emerging and less common modalities, even within the well-represented kinase complexes, Nittinger et al. reveal that cofolding models still struggle to place allosteric ligands into their allosteric pockets due to the over representation of orthosteric ligands in the PDB training data [22]. In summary, these studies expose the weaknesses of the current cofolding models, collectively suggesting directions for improvements of the next-generation cofolding models.

However, while these benchmarks successfully identify domain-specific challenges, they largely overlook a more fundamental requirement of modern SBDD: the capacity for ligand-modulated conformational reasoning. As detailed above, current evaluations remain predominantly ligand-centric, prioritizing the geometric placement of the small molecule through mostly pocket-aligned ligand Root Mean Square Deviation (RMSD) and overall global structural confidence in the protein fold. While these metrics are excellent starting points to verify basic structural integrity, they treat the protein receptor as a static scaffold rather than a dynamic landscape [23]. In practical drug discovery, identifying the correct global fold is necessary but not sufficient; successful lead optimization requires capturing the specific induced-fit conformational states dictated by ligand binding. Without a focused assessment of this conformational response, there remains a critical gap in understanding whether cofolding models can truly drive rational drug design.

The human kinase family presents an ideal paradigm to address this gap, given its strong representation in the PDB, its importance in therapeutic discovery [24–26], and the inherently dynamic nature of kinase conformational landscape [27–29]. Functioning as molecular switches, kinases toggle between active and inactive states via subtle rearrangements of the activation loop (DFG motif) and the *α*C-helix [30, 31]. More importantly, the utility of a kinase structure in drug discovery is defined by these specific conformational states, which dictate inhibitor binding kinetics and selectivity that helps distinguish Type I from Type II inhibitors [32]. Consequently, it is imperative and timely to evaluate the capacity of cofolding models to provide biophysically valid structural hypotheses, specifically by quantifying how well they capture the kinase conformational landscape and generalize across unseen ligand-induced states for drug discovery.

In this work, we introduce KinConfBench, a curated benchmark of 2,225 high-quality human kinase chains representing a total of 1,420 unique kinase systems (43 unique apo targets and 1,377 unique holo complexes), designed to shift the evaluation focus from ligand RMSD to protein-ligand conformational fidelity. To rigorously quantify these states, we employ the structural nomenclature developed by Modi and Dunbrack [31], labeling kinase chains with categorical structural features to capture specific active and inactive conformations. Using this framework, we evaluate three widely used open-source cofolding models—Boltz-2[15], Chai-1[13], and Protenix[16]—on their capability to recover experimentally observed kinase states in the presence of bound small-molecule ligands. The resulting metrics show that high confidence in the active site geometry does not ensure high confidence in predicting the kinase conformational states, especially when ligands perturb the kinase away from common templates of the global fold. We further document a critical generalization gap on kinase holo-apo pairs. When challenged with unseen holo states after the training cutoff, all three architectures increasingly predict the apo state, underscoring that the models are memorizing the apo-state baselines of the training set and poorly generalizing to true induced-fit outcomes. KinConfBench therefore foregrounds protein conformational correctness, not ligand RMSD alone, as the quantities that ultimately need to be ranked and interpreted to make progress in SBDD campaigns.

## Results

### The KinConfBench Conformational Benchmarks

To rigorously evaluate the capability of cofolding models to capture kinase conformational states, we curate KinConfBench, a high-quality benchmarking dataset derived from the RCSB PDB [7] as shown in Figure 1. To construct a comprehensive benchmarking dataset, we first identify 437 catalytically active human kinase domains and their corresponding UniProt [33] identifiers, following the classification established by Faezov and Dunbrack [34]. These UniProt IDs are then used to query the RCSB Protein Data Bank (PDB) [7] (snapshot 2025-08-01) to retrieve all associated structural entries. By prioritizing this UniProt-centric approach over broader sequence-similarity searches, we ensure the specific retrieval of human kinase domains while minimizing potential contamination from non-human homologs. This initial search yields a raw dataset of 7,763 unique PDB entries, each containing at least one relevant kinase chain.

**Figure 1:**
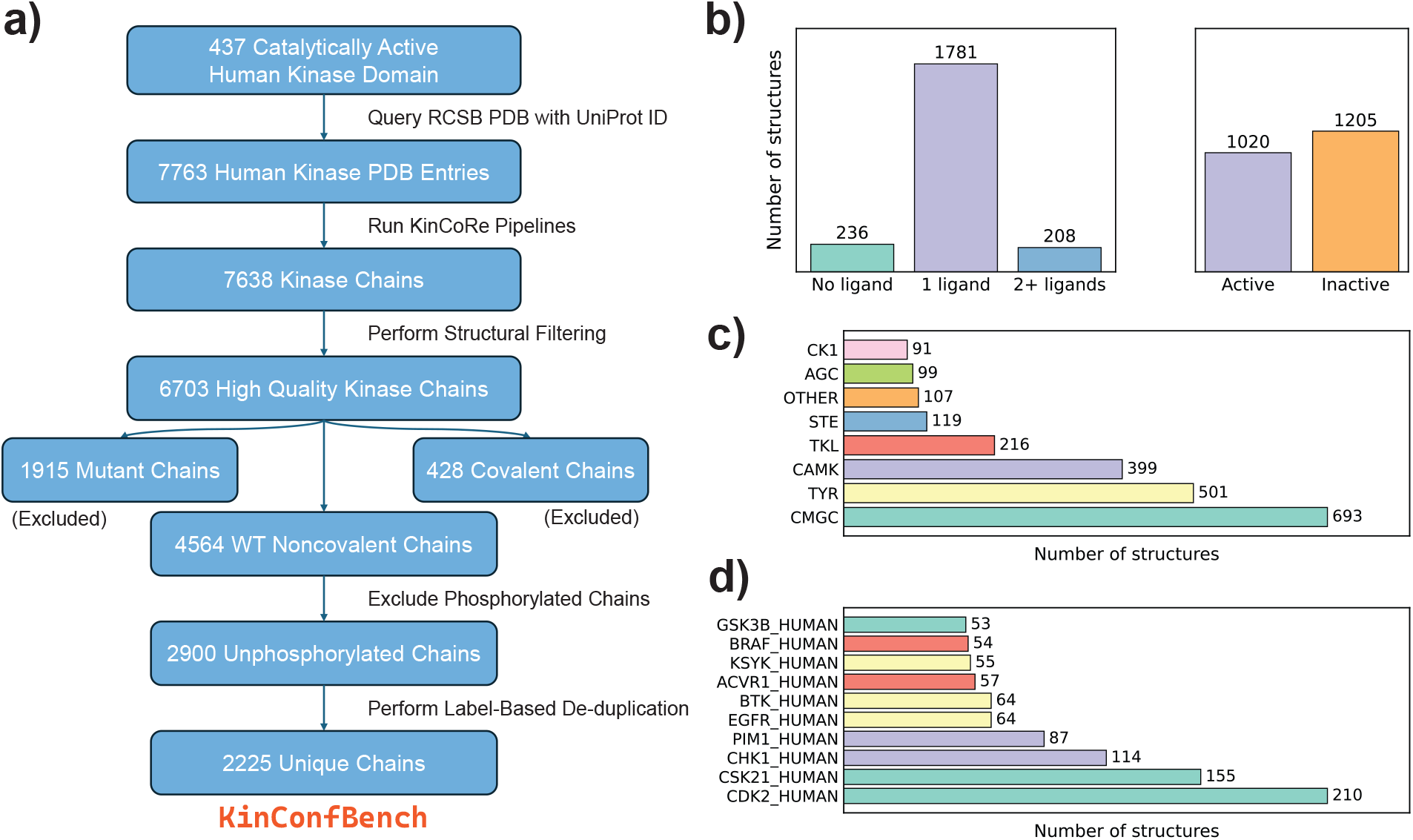
Overview of the KinConfBench curation pipeline and its data distribution. (a) Schematic of the curation workflow, detailing the sequence of filtering and labeling steps. (b) Distribution of ligand counts per kinase chain and their corresponding conformational activity. (c) Taxonomic distribution of kinase chains categorized by major kinase groups. Details for acronyms can be found in Supplementary Information. (d) Representation of the top 10 most frequent kinase genes in the dataset, colored by their kinase family.

To assign conformational labels to the selected kinase PDB entries, we process all chains from the identified entries using the KinCoRe software suite [35, 36]. KinCoRe categorizes kinase conformations by extracting and classifying key structural features into different conformational groups. There are a total of eight categorical labels: a spatial label that captures the orientation of the DFG motif; a dihedral label that captures the torsion angle distribution of the X-DFG motif; a helix label that captures the orientation of the *α*C-helix; a salt bridge label that captures the orientation of a conserved kinase salt bridge; N-terminus and C-terminus activation loop labels that capture the orientations of activation loops for both terminus; a spine label that captures the orientation of the kinase regulatory spine; and finally, the activity label that assigns whether the kinase structure is in an active or inactive form based on the previous labels. More details on the specific labeling format are provided in Supplementary Information.

Following the labeling process, we apply a stringent filtering protocol to exclude structures with missing atomic coordinates in critical functional regions that have “None” annotations in the KinCoRe output. Specifically, kinase chains are discarded if they have “None” values for either the spatial label or the helix label due to missing residues. This rigorous screening for structural motif completeness ensures that our benchmark is based on high-fidelity structural data, resulting in a refined set of 7,638 kinase chains.

To ensure stringent data quality control, we excluded chains with a resolution worse than 4.5 Å or those determined by NMR spectroscopy, following the protocol established by NeuralPlexer [10]. Furthermore, for genes encoding both catalytically active and pseudokinase domains, such as members of the Janus Kinase (JAK) family, we use the reference sequences of the active kinase domains extracted from UniProt to selectively retain only the catalytic units while excluding pseudokinase domains. This final filtering stage yields a high-quality set of 6,703 kinase chains.

The filtered dataset is partitioned into three primary categories: mutants (1,915 chains), covalent complexes (428 chains), and wild-type (WT) noncovalent complexes (4,564 chains). For this study, we focus exclusively on the unphosphorylated subset of the WT Noncovalent entries comprising 2,900 chains to benchmark co-folding model performances on canonical kinase sequences. To refine this subset, we remove trivial redundancies, defined as entries sharing identical sequences (and ligand SMILES, where applicable) alongside identical Kin-CoRe conformational labels. Nevertheless, we preserve meaningful structural diversity by retaining any entry that exhibited at least one unique KinCoRe label relative to others of the same condition. This logic is applied both to apo chains of the same gene and to holo complexes featuring the same protein-ligand combination, ensuring that our benchmark accounts for the multi-state nature of kinase domains. By prioritizing these distinct conformational states, we collect the final 2,225 structurally unique kinase chains for cofolding model benchmarking. This entire curation process is illustrated in Figure 1(a).

To investigate the general distribution of the selected unique kinase chains, we supply further analysis based on their gene properties and KinCoRe labels. As shown in Figure 1(b), the dataset is primarily composed of single-ligand holo complexes (∼80%), with apo structures and multi-ligand complexes each accounting for approximately 10%. This composition closely mirrors the broader distribution of kinases in the PDB, reflecting the data landscape encountered by co-folding models during training. Furthermore, while there appears to be a balanced global distribution between active and inactive states across the entire dataset, it is worth noting that this ratio can vary quite significantly at an individual gene level.

In terms of the kinase family distribution, we take the Manning classification system [37] and find that CMGC, TYR, CAMK, and TKL groups dominate the dataset, as these families represent areas of intense therapeutic interest[24–26, 38]. Indeed, the ten most frequently occurring genes in our dataset all belong to these four groups (Figure 1(c) and (d)). Further distributional metrics, including ligand and KinCoRe label frequencies, are detailed in Supplementary Figure S1. We observe a high prevalence of DFG-in and BLAminus labels, alongside a majority of Type I ligands that aligns with KinCoRe server results as shown in Supplementary Table S1. This observation indicates a persistent bias in the PDB toward holo complexes where the ligands occupy the ATP-binding pocket in catalytically active states, as the DFG-in/BLAminus configuration is a structural prerequisite for kinase activity [31]. After filtering for experimental resolution, sequence completeness, and distinct functional domain labels, the final curated dataset of 2,225 kinase chains is comprised of 236 unique apo chains representing 43 unique gene targets, and 1,989 unique holo chains with 1,377 involving unique protein-ligand pairs, ensuring broad coverage of the human kinome.

### Evaluation of Boltz-2, Chai-1, and Protenix against KinConfBench

Protein kinases are highly dynamic molecular machines that transition between multiple active and inactive isoforms to perform their function[27–29]. Molecular effectors such as small drug molecules introduce new interactions that can shift the thermodynamic equilibrium among these states. For instance, Type-II inhibitors specifically recognize and stabilize the inactive “DFG-out” conformation by binding into an adjacent, allosteric hydrophobic pocket [39, 40]. By establishing favorable new bonds within this newly exposed site, the ligand significantly lowers the potential energy minimum of the inactive state relative to the active one [41], ad the kinase becomes thermodynamically trapped in this non-functional shape, effectively halting its catalytic activity. Therefore in the following sections we evaluate not only the ligand-centric geometric performance of the cofolding models, but establish the necessity of providing conformational selection metrics to enable drug selectivity for kinase targets.

### From geometric filters to functional conformations

Valid and conformationally diverse protein folded states is a fundamental prerequisite for the meaningful use of cofolding models. KinCoRe assigns each kinase protein structure to valid spatial and dihedral label combinations (36 theoretical, 12 observed in the PDB; Supplementary Table S1) [31]. By applying the KinCoRe labels to every prediction, we establish a “ground truth” based on the categorical orientation of the DFG motif and the *α*C-helix, providing a more functionally relevant metric than standard backbone RMSD. Mapping the inference results against the established kinase clusters reveals that all cofolding models generate protein structures falling exclusively within valid conformational categories as shown in Supplementary Table S2, no “forbidden” or physically unreasonable protein geometries appear, confirming that all tested cofolding models possess the baseline biophysical reasoning required to produce structurally plausible kinase conformations.

Beyond biophysical validity of the protein fold, a cofolding model must place the ligand correctly into the pocket, which we characterize using three structural metrics. Global fold quality, lDDT-C*α*, summarizes C*α* agreement of the predicted complex to the experimental structure; higher values indicate a closer global backbone match. Pocket-focused geometry, lDDT-PLI, emphasizes the agreement of the binding site and ligand-adjacent protein atoms compared to experimental values. Finally, ligand heavy-atom RMSD indicates agreement of the modeled ligand pose with the crystal pose. We define a geometrically successful prediction as one with global fold lDDT-C*α* ≥ 0.7, binding pocket lDDT-PLI ≥ 0.8, and ligand RMSD < 2.0Å following the definition of previous works [2, 17]. Out of 1420 protein-ligand kinase systems, geometric filtering resulted in only 950 kinase where all cofolding models produce at least one geometrically valid pose out of 20 samples for downstream analysis. The broad geometric distribution of these successful predictions for each cofolding model across kinase systems is detailed in Supplementary Figure S2.

However, a geometrically successful binding pose does not ensure that the cofolding model has captured the correct induced conformational state. Specifically, a prediction is only considered “correct” if all eight KinCoRe labels match those of the experimental structures. Figure 2(a) shows the relation of the three geometric metrics (lDDT-C*α*, lDDT-PLI, and ligand RMSD) to KinCoRe label correctness across all models. While correct conformational assignments skew better across the geometric metrics, the overlap with incorrect functional assignment remains large. The side-by-side comparison among the cofolding models highlights that mislabeled states can still achieve strong ligand-centric structural scores around the binding pocket, indicating that geometry alone is an incomplete proxy for robust cofolding prediction.

**Figure 2:**
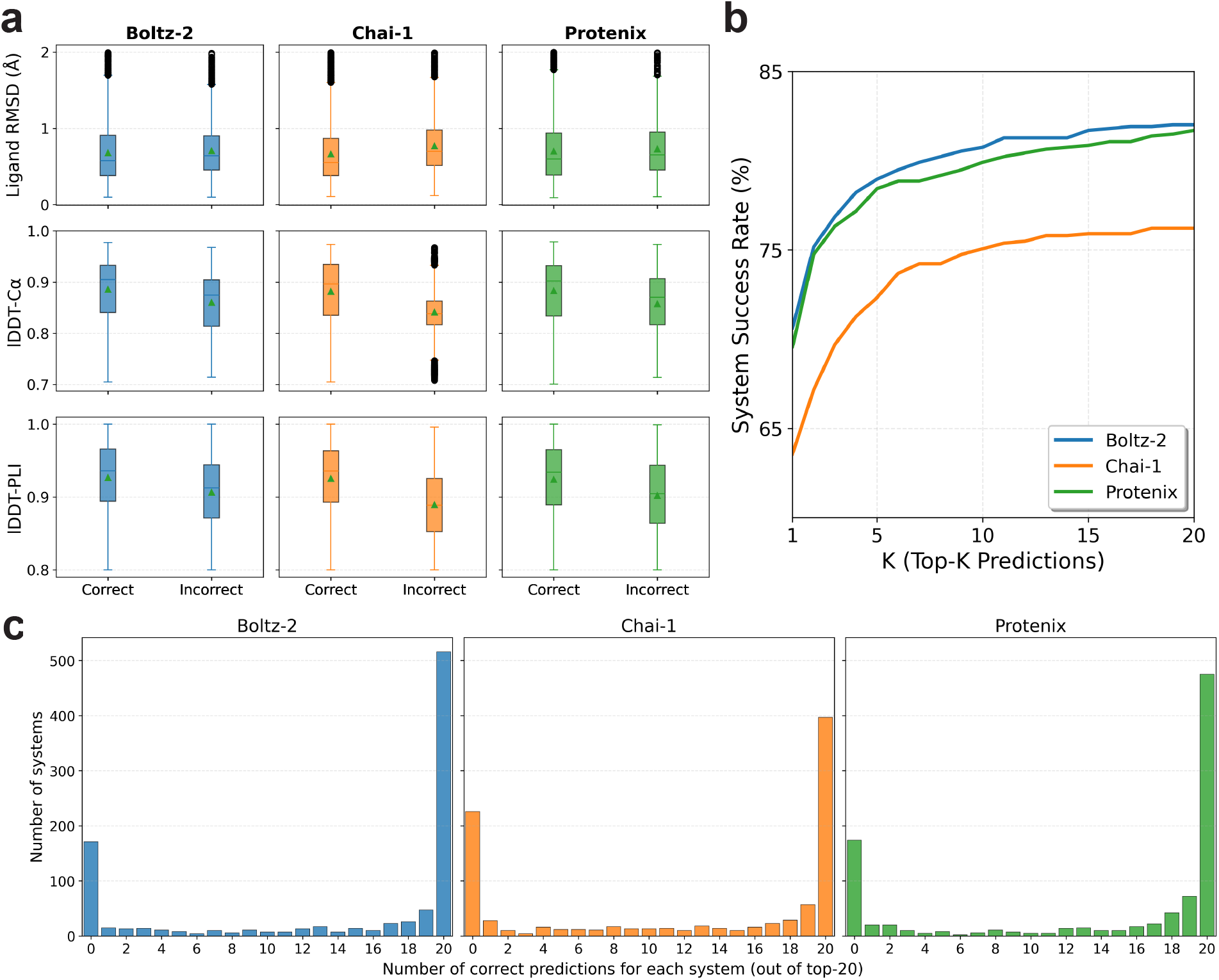
Correct and incorrect KinCoRe classification for all three cofolding model predictions passing geometric filters. (a) Cofolding model performance on geometric predictions (lDDT-C*α* ≥ 0.7, lDDT-PLI ≥ 0.8, ligand RMSD < 2.0 Å) comparing samples with all eight KinCoRe labels being correct versus incorrect. To account for intrinsic structural flexibility where a single experimental entry may capture multiple conformational states, we consider a prediction successful if it matches any valid set of ground truth labels associated with that protein-ligand complex. (b) Success rate of each cofolding model versus a correct ranked prediction among the top-*K* by confidence. For each target, we evaluate an ensemble of *N* = 20 generated structures, ranked by their pLDDT confidence scores. (c) Histogram of the count of top-20 predictions that match all KinCoRe labels of their corresponding experimental structures. The *x*-axis is the number of correct predictions in the ranked ensemble (0-20), and the *y*-axis is how many of the 950 kinase systems fall in each bin. Panels share a common vertical scale to contrast how frequently each architecture saturates the ensemble with correct states versus how often it fails to identify a single exact match.

Figure 2(b) shows each co-folding models’ success versus *K*, defined as any correct functional state among the top-*K* by confidence ranking for each cofolding model. Both Boltz-2 and Protenix lead at Top-1 compared to Chai-1, while moving from Top-1 to Top-20 adds roughly 11-13 percentage points for each model (Boltz-2 +11.4; Chai-1 +12.6; Protenix +12.1), indicating that additional samples sometimes recover the correct conformational state even when the highest-confidence pose is wrong (see also Supplementary Table S3). However, Figure 2(c) shows that, despite the increase in accuracy, the co-folding model predictions are largely bimodal. This mostly all-right or all-wrong pattern suggests ensemble mode collapse rather than graded uncertainty within the twenty samples.

Supplementary Table 4 contains the 110 kinases where all three cofolding models fail to generate the correct conformation, providing an important benchmark for future cofolding model improvements. Figure 3 highlights MAP4K1/YK1 (PDB ID: 7M0M) [42] as a representative hard case where all cofolding model predictions fail to obtain an induced-fit to the correct conformational state. This observation shows that even when pocket metrics and ligand poses look favorable, Boltz-2, Chai-1, and Protenix often jointly settle into an alternate spatial and dihedral configuration that disagrees with the experimentally observed loop excursion, underscoring why ligand RMSD alone can mask protein-state errors.

**Figure 3:**
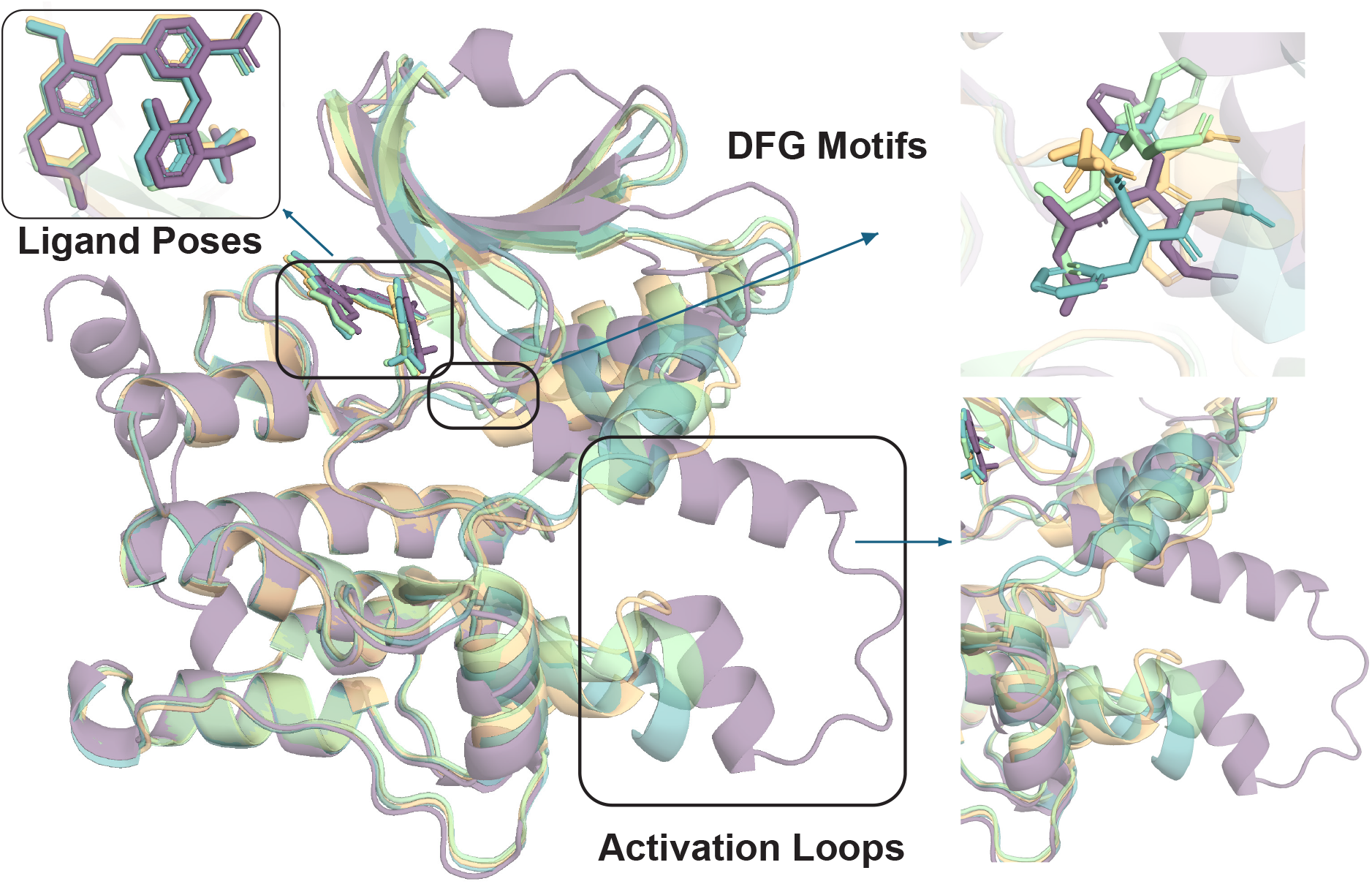
Induced-fit failure modes for MAP4K1 (7M0M). Superposition of the experimental Type-I diaminopyrimidine carboxamide complex (purple) with representative Boltz-2 (green), Chai-1 (orange), and Protenix (blue) predictions evaluated in KinConfBench. The experimental structure adopts DFG-in, BLAminus packing with an inward *α*C-helix and an extended, outward-biased activation loop. **(Top right)** DFG region; **(bottom right)** activation loop. Although ligand placements are perfect across models, the three evaluated cofolding predictions miss the cooperative loop sampled in the PDB, collapsing instead toward structure that is conformationally mislabeled relative to 7M0M.

### Ensemble Diversity and Resolution of Complex Targets

KinCoRe labels establish whether a model satisfies the mechanistic requirements of a kinase state such as satisfying the DFG and *α*C-helix motifs, but categorical success does not capture the nuances of “intra-basin exploration.” Hence, we extend our analysis beyond discrete accuracy to measure the structural dispersion within ensembles that are already deemed functionally correct. To quantify conformational diversity we use some of the same geometric descriptors that define the KinCoRe labels as well as some additional metrics, but on a continuous scale for distances and dihedral angles as described in Supplementary Information. For this benchmark we focus on the 509 systems where Boltz-2, Chai-1, and Protenix each have at least ten all-correct structures according to KinCoRe in the top-20, so every cofolding method contributes a comparably populated success set based on classification; Supplementary Figure S3 summarizes how these targets distribute across the Top-K benchmark.

Figure 4 shows the standard deviation in distance and torsion angle metrics around their average values across the 20 member ensembles for all three cofolding models. While Supplementary Figure S4 finds that all cofolding models capture the correct average distance and angle values compared to experiment, all models produce very small structural deviations of ∼0.1 Å for the distance metrics and less than 5° average angle exploration except for Chai-1’s performance on Asp *χ*_2_. To place these results into context, root mean square fluctuations of ∼1-2 Å and backbone or side chain dihedral angle ranges of 5° to 30° define the statistical fluctuations of stable protein conformations [43, 44], while significantly higher values would be needed to indicate transitions between distinct conformational states. Hence while the Chai-1 model shows greater explorations within the correct structural basin compared to Boltz-2 and Protenix, on an absolute scale the conformational diversity is still quite limited, in line with previous ligand-centric analysis of cofolding models[17].

**Figure 4:**
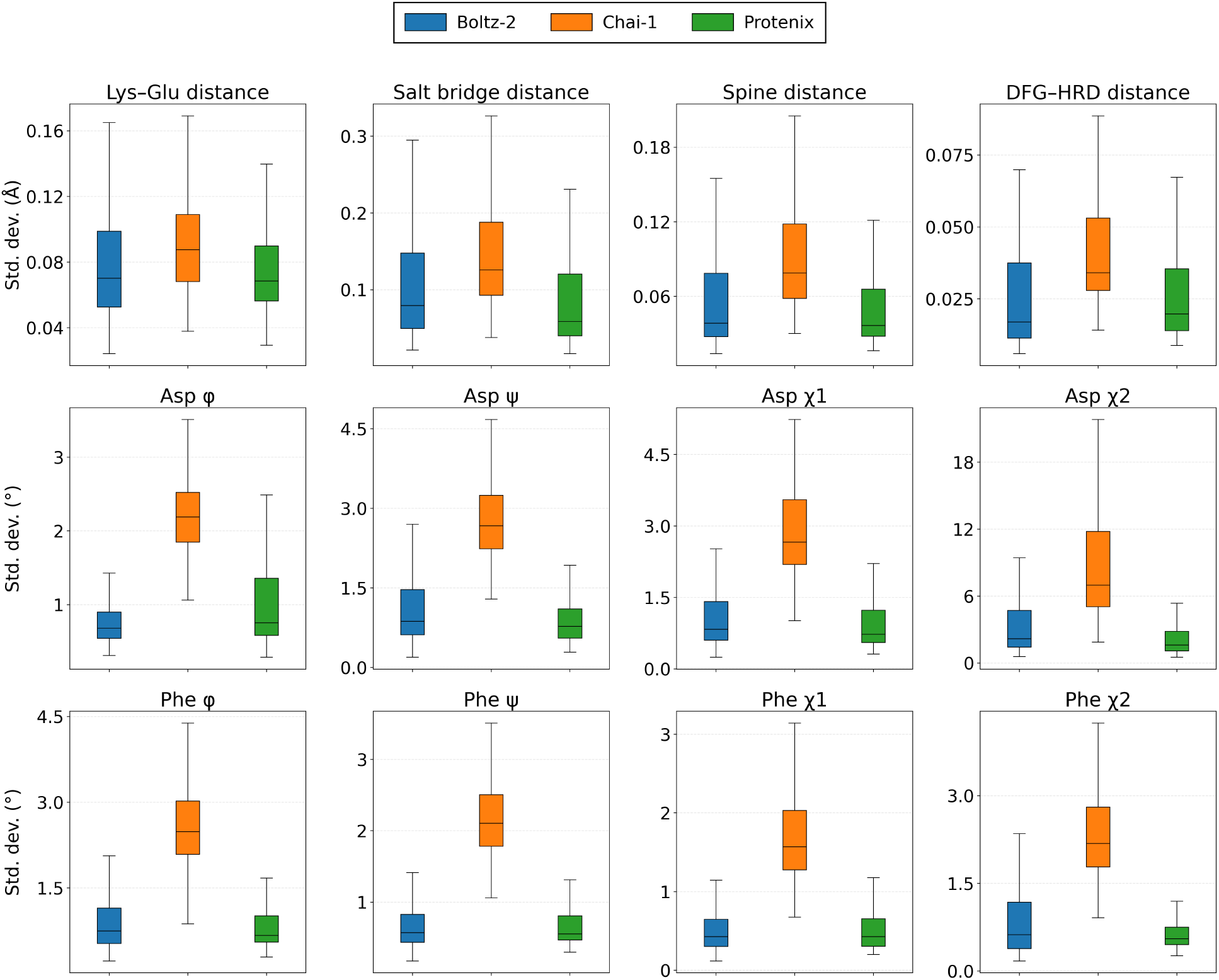
Distance and torsion angle ensemble diversity among correct KinCoRe predictions across cofolding models. For each system in the diversity subset from KinConBench, we retain only cofolding predictions that are all-label correct, then measure the distance and dihedral angle standard deviations across those correct poses. Overall, all cofolding models have little structural diversity. Panels compare Boltz-2 (blue), Chai-1 (orange), and Protenix (green).

### Benchmarking Apo-to-Holo Conformational Shift

In structure-based drug design, ligand binding represents a thermodynamic perturbation of the protein’s native conformational landscape. We conceptualize the apo (ligand-free) distribution as a cofolding model’s learned default energy landscape, whereas the introduced ligand acts as a physical modulator that distorts this baseline to stabilize a distinct holo conformation. If a cofolding model learns the underlying physics of molecular recognition, it successfully predicts this specific ligand-induced apo-to-holo conformational shift. Conversely, if a cofolding model relies heavily on memorization it will fail to generalize to an apo-to-holo conformational transition for novel ligand effectors, effectively “drifting” back to the unperturbed, default apo state.

To test this hypothesis, we perform a comparative analysis against the ground-truth holo and apo state pairs. We extract kinase systems from KinConfBench that possess resolved structures for both states, restricting our analysis to systems where the ligand induces a distinct structural change defined as a different set of KinCoRe functional labels compared to ligand-free structures. For each cofolding model, we define training and test sets based on a time split relevant to each model: a holo ligand combination counts as part of the test set if it first appears in the PDB only after that method’s training-data deposition cutoff: Boltz-2 (2023-06-01); Chai-1 (2021-01-12); Protenix (2021-09-30)). All apo-state structures are found in the training data for all cofolding models.

For each of the kinase system pairs, we generate an ensemble of 20 structures using the holo sequence-ligand as input for each cofolding model, and we classify their predictions according to:

- **Holo State:** At least one structure in the generated ensemble exactly matches the KinCoRe labels of the ground-truth holo reference and 0 matches the KinCoRe labels of the ground-truth apo reference. This metric quantifies the model’s ability to correctly predict the ligand-induced state.
- **Apo Drift:** At least one structure in the generated ensemble exactly matches the KinCoRe labels of the ground-truth apo reference and 0 matches the KinCoRe labels of the ground-truth holo reference. This metric quantifies the tendency of the model to revert to the memorized apo state despite the presence of the ligand.
- **Dual Match:** At least one structure of the ensemble matches the KinCoRe labels of the ground-truth holo reference, and another matches the ground-truth apo reference. This metric quantifies the possibility that the holo-state could be recovered with more sampling or an effective ranking, but otherwise is also evidence of apo-drift.
- **None:** No structure in the generated ensemble exactly matches the KinCoRe labels of either the ground-truth holo reference or the ground-truth apo reference.

A robustly generalizing cofolding model should keep a large Holo-State fraction on both the training and testing data, whereas memorization shifts toward Apo-only or dual-match states predictions. As seen in Figure 5a, within the training set, the Holo-State prediction represents roughly 40% of the predictions for Boltz-2 and Protenix and around 30% for Chai-1, but all three models display nearly 40% Apo Drift within the training set. Moreover, when comparing to the test set distribution in Figure 5b against the training set, the stacked profiles further shift toward Apo-only and Dual-Match segments relative to the training systems—most clearly for Boltz-2 and Protenix—consistent with apo-drift when confronted with new holo states unsupported in the training statistics.

**Figure 5:**
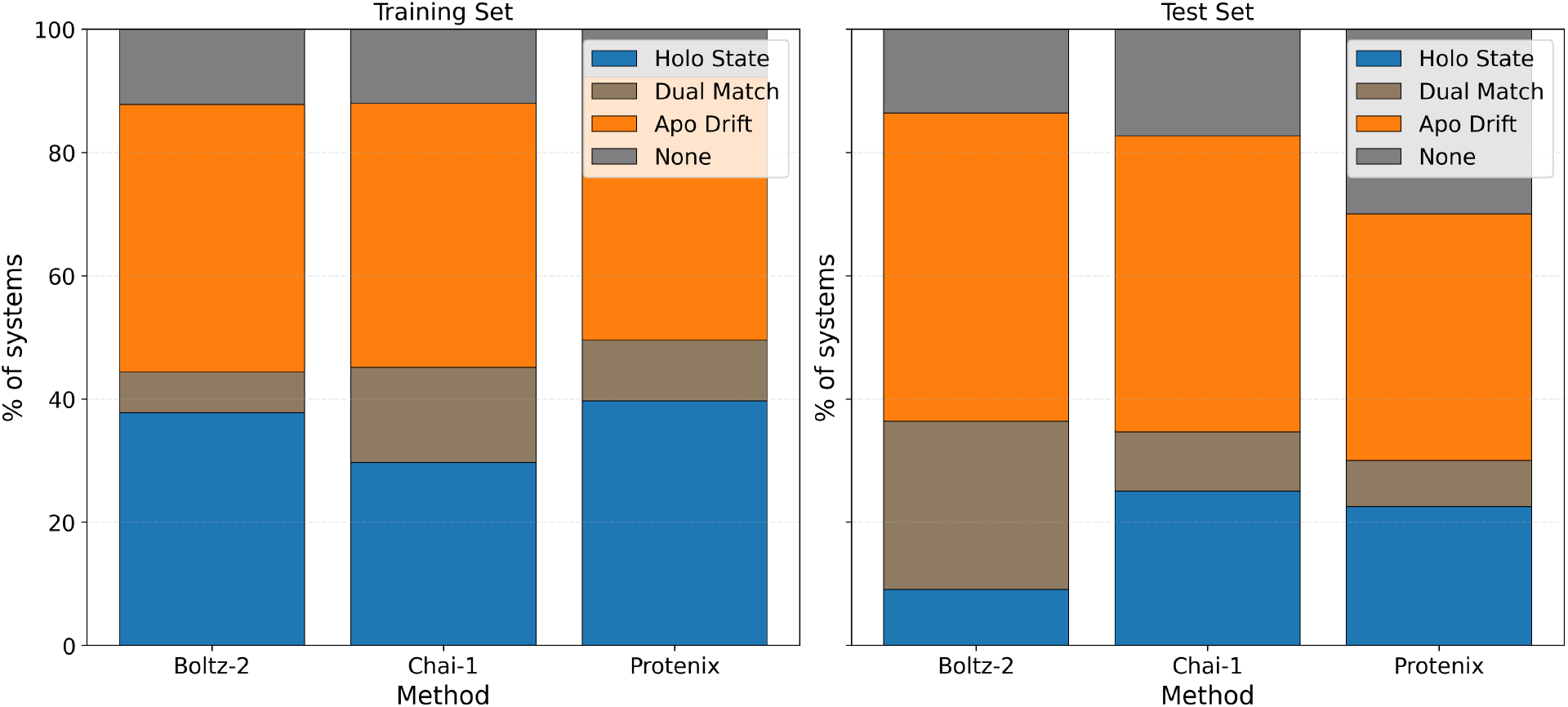
Generalization and apo-drift across co-folding models. Stacked fraction of systems in each holo/apo outcome category on holo→apo inference pairs for Boltz-2, Chai-1, and Protenix. Sample sizes per split (Train / Test systems): Boltz-2 212 / 22; Chai-1 182 / 52; Protenix 194 / 40. Categories (bottom to top in stack): Holo only; Dual Match; Apo only; None.

## Discussion

Our benchmarking of three popular cofolding models highlights a tension between confident geometric poses and biophysically faithful kinase conformational states. Each architecture attains strong pocket-centric metrics on large portions of KinConfBench, yet functional Kin-CoRe agreement lags behind what raw lDDT or RMSD might suggest. Framing success purely through the highest-confidence structure therefore obscures a clinically relevant failure mode: models can “look” bound while adopting the wrong regulatory layout for the ligand at hand.

Ensemble behavior compounds the issue for all cofolding models. When additional samples are drawn, Top-20 accuracy gains indicate that correct kinase conformational states sometimes appear deeper in the ranking, but per-target histograms remain dominated by all-or-nothing outcomes rather than graded uncertainty. None of the three cofolding models eliminates the need for external checks when structure-activity-relationships involving induced fit critical motions are required, as in MAP4K1/7M0M, where several models conspire on an alternate DFG-in basin. This likely arises because even when a cofolding model correctly predicts the KinCoRe classification, structural variety is seen to be well-below that expected for thermalized statistical fluctuations needed to create induced-fit configurations.

The holo/apo paired analysis sharpens this diagnosis. Memorization of apo kinase states is shared across Boltz-2, Chai-1, and Protenix rather than unique to one model. After the training cutoff (specific to each model), we find that the holo state prediction rates fall and apo drift rises for each architecture. Closing this gap likely demands both better generative breadth for rare induced-fit basins and confidence objectives that reward state discrimination, not only ligand-protein pocket aligned metrics.

At its core, drug discovery requires conformational state correctness rather than solely fold correctness or memorization of training instances. Predicting the general architecture of a kinase is biologically insufficient if the specific, ligand-induced functional conformation is wrong. KinConfBench is meant to keep that standard explicit as cofolding models continue to mature. Looking forward, the scope of kinase benchmarking must expand to encompass the intricate regulatory mechanisms that define this protein family in their real biological environment. Our current work focuses on the interaction between wild-type domains and non-covalent inhibitors, yet kinase signaling is tightly modulated by post-translational modifications (PTMs) and somatic mutations. Future benchmarks should rigorously assess whether cofolding models can predict how phosphorylation at specific regulatory sites or drug-resistant mutations alter the conformational ensemble. Furthermore, the rise of covalent inhibitors in oncology necessitates a focused evaluation of how cofolding models handle the shift in protein context when a covalent bond is formed, as covalent ligands typically bring more unforseen perturbations than simple non-covalent docking.

Finally, the ultimate test of a cofolding model’s biophysical reasoning will be its ability to simulate competitive dynamics. In a physiological environment, kinases are rarely isolated with just a single ligand. For Type-I kinase inhibitors, they constantly exist in a competitive environment where inhibitors must displace ATP or compete with other endogenous substrates. Future generations of benchmarks should evaluate the ability of models to handle multi-ligand scenarios, probing whether the presence of competing binders drives the protein into distinct conformational equilibria. Moving beyond static snapshots to dynamic, competitive scenarios will be essential for evolving these tools from structural prediction engines into true computational assays for drug discovery.

## Materials and Methods

### Details of and Inference from the Cofolding Models

When performing inferences for Boltz-2 [15], Chai-1 [13], and Protenix [16], we adhere to the official inference pipelines for each method to represent standard usage scenarios. Multiple Sequence Alignments (MSAs) are generated and cached with each tool’s default databases and pairing strategies. For Boltz-2, Chai-1, and Protenix, the ColabFold server [3] is used for MSA generation, while sequence pairing follows the defaults of the respective model repositories. Input sequences are restricted to the canonical kinase domain boundaries defined by UniProt, so that the benchmark stresses domain-level cofolding behavior.

Inferences are performed on NVIDIA A100 GPUs across the 1,420 unique configuration pairs (43 apo + 1,377 holo) defined during curation. Post-inference, structures are passed through the Dunbrack KinCoRe annotation pipeline; targets that failed annotation because of malformed outputs or missing motifs are excluded. To ensure robust sampling of the conformational landscape, we generate *N* = 20 cofolding samples per target for each inference run. Output structures are saved in mmCIF format to ensure compatibility with the KinCoRe annotation pipeline.

## Supplementary Information

Details of additional KinConfBench analysis of all cofolding models and the KinCoRe pipelines and the 110 systems where all cofolding predictions fail to generate the correct conformation as the ground truth are provided in the Supplementary Information.

## Data and Code Availability

All the codes and data are available on GitHub: https://github.com/THGLab/KinConfBench

## Acknowledgments

We thank Mia Rosenfeld for contributions to project conception, experimental design, and narrative framing. We also thank Matthew Welborn, Aleksander Durumeric, and Michael Irvin for helpful suggestions. This work was supported by National Institute of Allergy and Infectious Disease grant U19-AI171954. This research used computational resources of the National Energy Research Scientific Computing, a DOE Office of Science User Facility supported by the Office of Science of the U.S. Department of Energy under Contract No. DE-AC02-05CH11231.

## Author Contributions

KS conceived the project and wrote the code and performed all computational experiments. KS and THG did all analysis and wrote the paper.

## References

[1] Baek M, DiMaio F, Anishchenko I, Dauparas J, Ovchinnikov S, Lee GR, et al. Accurate prediction of protein structures and interactions using a three-track neural network. Science. 2021 Aug;373(6557):871–6. Available from: https://www.science.org/doi/10.1126/science.abj8754.

[2] Jumper J, Evans R, Pritzel A, Green T, Figurnov M, Ronneberger O, et al. Highly accurate protein structure prediction with AlphaFold. Nature. 2021 Aug;596(7873):583–9. Available from: https://www.nature.com/articles/s41586-021-03819-2.

[3] Mirdita M, Schütze K, Moriwaki Y, Heo L, Ovchinnikov S, Steinegger M. ColabFold: making protein folding accessible to all. Nature Methods. 2022 Jun;19(6):679–82. Available from: https://www.nature.com/articles/s41592-022-01488-1.

[4] Lin Z, Akin H, Rao R, Hie B, Zhu Z, Lu W, et al. Evolutionary-scale prediction of atomic-level protein structure with a language model. Science. 2023 Mar;379(6637):1123–30. Available from: https://www.science.org/doi/10.1126/science.ade2574.

[5] Fang X, Wang F, Liu L, He J, Lin D, Xiang Y, et al. A method for multiple-sequence-alignment-free protein structure prediction using a protein language model. Nature Machine Intelligence. 2023 Oct;5(10):1087–96. Available from: https://www.nature.com/articles/s42256-023-00721-6.

[6] Ahdritz G, Bouatta N, Floristean C, Kadyan S, Xia Q, Gerecke W, et al. OpenFold: retraining AlphaFold2 yields new insights into its learning mechanisms and capacity for generalization. Nature Methods. 2024 Aug;21(8):1514–24. Available from: https://www.nature.com/articles/s41592-024-02272-z.

[7] Berman HM. The Protein Data Bank. Nucleic Acids Research. 2000 Jan;28(1):235–42. Available from: https://academic.oup.com/nar/article-lookup/doi/10.1093/nar/28.1.235.

[8] Evans R, O’Neill M, Pritzel A, Antropova N, Senior A, Green T, et al. Protein complex prediction with AlphaFold-Multimer; 2021. Available from: http://biorxiv.org/lookup/doi/10.1101/2021.10.04.463034.

[9] Gao M, Nakajima An D, Parks JM, Skolnick J. AF2Complex predicts direct physical interactions in multimeric proteins with deep learning. Nature Communications. 2022 Apr;13(1):1744. Available from: https://www.nature.com/articles/s41467-022-29394-2.

[10] Qiao Z, Nie W, Vahdat A, Miller TF, Anandkumar A. State-specific protein–ligand complex structure prediction with a multiscale deep generative model. Nature Machine Intelligence. 2024 Feb;6(2):195–208. Available from: https://www.nature.com/articles/s42256-024-00792-z.

[11] Abramson J, Adler J, Dunger J, Evans R, Green T, Pritzel A, et al. Accurate structure prediction of biomolecular interactions with AlphaFold 3. Nature. 2024 Jun;630(8016):493–500. Available from: https://www.nature.com/articles/s41586-024-07487-w.

[12] Krishna R, Wang J, Ahern W, Sturmfels P, Venkatesh P, Kalvet I, et al. Generalized biomolecular modeling and design with RoseTTAFold All-Atom. Science. 2024 Apr;384(6693):eadl2528. Available from: https://www.science.org/doi/10.1126/science.adl2528.

[13] Chai Discovery, Boitreaud J, Dent J, McPartlon M, Meier J, Reis V, et al. Chai-1: Decoding the molecular interactions of life; 2024. Available from: http://biorxiv.org/lookup/doi/10.1101/2024.10.10.615955.

[14] Wohlwend J, Corso G, Passaro S, Getz N, Reveiz M, Leidal K, et al. Boltz-1: Democratizing Biomolecular Interaction Modeling. bioRxiv. 2024.

[15] Passaro S, Corso G, Wohlwend J, Reveiz M, Thaler S, Somnath VR, et al. Boltz-2: Towards Accurate and Efficient Binding Affinity Prediction. bioRxiv. 2025.

[16] ByteDance AML AI4Science Team, Chen X, Zhang Y, Lu C, Ma W, Guan J, et al. Protenix - Advancing Structure Prediction Through a Comprehensive AlphaFold3 Reproduction; 2025. Available from: http://biorxiv.org/lookup/doi/10.1101/2025.01.08.631967.

[17] Škrinjar P, Eberhardt J, Tauriello G, Schwede T, Durairaj J. Have protein-ligand cofolding methods moved beyond memorisation?; 2025. Available from: http://biorxiv.org/lookup/doi/10.1101/2025.02.03.636309.

[18] Buttenschoen M, M Morris G, M Deane C. PoseBusters: AI-based docking methods fail to generate physically valid poses or generalise to novel sequences. Chemical Science. 2024;15(9):3130–9. Available from: https://pubs.rsc.org/en/content/articlelanding/2024/sc/d3sc04185a.

[19] Zhang Th, Zhu Jt, Huang Zx, Xie J, Pei Jf, Lai Lh. Benchmarking co-folding methods to predict the structures of covalent protein–ligand complexes. Acta Pharmacologica Sinica. 2026 Jan:1–11. Available from: https://www.nature.com/articles/s41401-025-01721-5.

[20] Dunlop N, Erazo F, Jalalypour F, Mercado R. Predicting PROTAC-mediated ternary complexes with AlphaFold3 and Boltz-1. Digital Discovery. 2025;4(12):3782–809. Available from: https://pubs.rsc.org/en/content/articlelanding/2025/dd/d5dd00300h.

[21] Liao Y, Zhu J, Xie J, Lai L, Pei J. Benchmarking Cofolding Methods for Molecular Glue Ternary Structure Prediction. Journal of Chemical Information and Modeling. 2025 Oct;65(20):11136–48. Available from: https://pubs.acs.org/doi/10.1021/acs.jcim.5c01860.

[22] Nittinger E, Yoluk O, Tibo A, Olanders G, Tyrchan C. Co-folding, the future of docking – prediction of allosteric and orthosteric ligands. Artificial Intelligence in the Life Sciences. 2025 Dec;8:100136. Available from: https://www.sciencedirect.com/science/article/pii/S2667318525000121.

[23] Wei H, McCammon JA. Structure and dynamics in drug discovery. npj Drug Discovery. 2024 Nov;1(1):1. Available from: https://www.nature.com/articles/s44386-024-00001-2.

[24] Wang Yy, Zhao R, Zhe H. The emerging role of CaMKII in cancer. Oncotarget. 2015 Apr;6(14):11725–34. Available from: https://www.ncbi.nlm.nih.gov/pmc/articles/PMC4494900/.

[25] Zhang M, Zhang L, Hei R, Li X, Cai H, Wu X, et al. CDK inhibitors in cancer therapy, an overview of recent development. American Journal of Cancer Research. 2021 May;11(5):1913–35. Available from: https://www.ncbi.nlm.nih.gov/pmc/articles/PMC8167670/.

[26] Pottier C, Fresnais M, Gilon M, Jérusalem G, Longuespée R, Sounni NE. Tyrosine Kinase Inhibitors in Cancer: Breakthrough and Challenges of Targeted Therapy. Cancers. 2020 Mar;12(3):731. Available from: https://www.ncbi.nlm.nih.gov/pmc/articles/PMC7140093/.

[27] McClendon CL, Kornev AP, Gilson MK, Taylor SS. Dynamic architecture of a protein kinase. Proceedings of the National Academy of Sciences of the United States of America. 2014 Oct;111(43):E4623–31.

[28] Taylor SS, Kornev AP. Protein kinases: evolution of dynamic regulatory proteins. Trends in Biochemical Sciences. 2011 Feb;36(2):65–77.

[29] Taylor SS, Wu J, Bruystens JGH, Del Rio JC, Lu TW, Kornev AP, et al. From structure to the dynamic regulation of a molecular switch: A journey over 3 decades. The Journal of Biological Chemistry. 2021;296:100746.

[30] Möbitz H. The ABC of protein kinase conformations. Biochimica Et Biophysica Acta. 2015 Oct;1854(10 Pt B):1555–66.

[31] Modi V, Dunbrack RL. Defining a new nomenclature for the structures of active and inactive kinases. Proceedings of the National Academy of Sciences. 2019 Apr;116(14):6818–27. Available from: https://pnas.org/doi/full/10.1073/pnas.1814279116.

[32] Roskoski R. Classification of small molecule protein kinase inhibitors based upon the structures of their drug-enzyme complexes. Pharmacological Research. 2016 Jan;103:26–48. Available from: https://www.sciencedirect.com/science/article/pii/S1043661815301298.

[33] The UniProt Consortium, Bateman A, Martin MJ, Orchard S, Magrane M, Adesina A, et al. UniProt: the Universal Protein Knowledgebase in 2025. Nucleic Acids Research. 2025 Jan;53(D1):D609–17. Available from: https://academic.oup.com/nar/article/53/D1/D609/7902999.

[34] Faezov B, Dunbrack RL. AlphaFold2 models of the active form of all 437 catalytically competent human protein kinase domains; 2023. Available from: http://biorxiv.org/lookup/doi/10.1101/2023.07.21.550125.

[35] Modi V, Dunbrack RL. Kincore: a web resource for structural classification of protein kinases and their inhibitors. Nucleic Acids Research. 2022 Jan;50(D1):D654–64. Available from: https://academic.oup.com/nar/article/50/D1/D654/6395339.

[36] Faezov B, Dunbrack RL. AlphaFold2 models of the active form of all 437 catalytically competent human protein kinase domains; 2023. Available from: http://biorxiv.org/lookup/doi/10.1101/2023.07.21.550125.

[37] Manning G, Whyte DB, Martinez R, Hunter T, Sudarsanam S. The Protein Kinase Complement of the Human Genome. Science. 2002 Dec;298(5600):1912–34. Available from: https://www.science.org/doi/10.1126/science.1075762.

[38] Śmiech M, Leszczynśki P, Kono H, Wardell C, Taniguchi H. Emerging BRAF Mutations in Cancer Progression and Their Possible Effects on Transcriptional Networks. Genes. 2020 Nov;11(11):1342. Available from: https://www.ncbi.nlm.nih.gov/pmc/articles/PMC7697059/.

[39] Kufareva I, Abagyan R. Type-II Kinase Inhibitor Docking, Screening, and Profiling Using Modified Structures of Active Kinase States. Journal of Medicinal Chemistry. 2008 Dec;51(24):7921–32. Available from: https://pubs.acs.org/doi/10.1021/jm8010299.

[40] Vijayan RSK, He P, Modi V, Duong-Ly KC, Ma H, Peterson JR, et al. Conformational Analysis of the DFG-Out Kinase Motif and Biochemical Profiling of Structurally Validated Type II Inhibitors. Journal of Medicinal Chemistry. 2015 Jan;58(1):466–79. Available from: https://pubs.acs.org/doi/10.1021/jm501603h.

[41] Zhao Z, Wu H, Wang L, Liu Y, Knapp S, Liu Q, et al. Exploration of Type II Binding Mode: A Privileged Approach for Kinase Inhibitor Focused Drug Discovery? ACS Chemical Biology. 2014 Jun;9(6):1230–41. Available from: https://pubs.acs.org/doi/10.1021/cb500129t.

[42] Lesburg CA. HPK1 IN COMPLEX WITH COMPOUND 1: 7m0m; 2021. Available from: https://www.wwpdb.org/pdb?id=pdb_00007m0m.

[43] Kalaivani R, Narwani TJ, De Brevern AG, Srinivasan N. Long-range molecular dynamics show that inactive forms of Protein Kinase A are more dynamic than active forms. Protein Science. 2019 Mar;28(3):543–60. Available from: https://onlinelibrary.wiley.com/doi/10.1002/pro.3556.

[44] Shapovalov MV, Dunbrack RL. A smoothed backbone-dependent rotamer library for proteins derived from adaptive kernel density estimates and regressions. Structure. 2011;19(6):844–58.

