## Supplementary material for "KinConfBench: A Curated Benchmark for Cofolding Models on Kinase Conformational States": SI

**S1 Additional Methodology Details** **2**

**S2 Supplementary Tables** **4**

**S3 Supplementary Figures** **6**

### S1 Additional Methodology Details

**Kinase Group Classifications.** To categorize the structural and functional diversity of the kinome, we utilize the standard Manning classification system [1]. The primary groups are defined as follows.

- **AGC**: Proteins including PKA, PKG, and PKC families.
- **CAMK**: Calcium/calmodulin-dependent protein kinases.
- **CK1**: Casein kinase 1 and closely related isoforms.
- **CMGC**: Proteins including CDK, MAPK, GSK3, and CLK families.
- **STE**: Homologs of yeast Sterile 7, Sterile 11, and Sterile 20 kinases.
- **TYR**: Tyrosine kinases.
- **TKL**: Tyrosine kinase–like proteins that are serine-threonine kinases.
- **OTHER**: Other kinases that don't fall into above classification.

**Details of KinCoRe Annotations.** Introduced by Modi and Dunbrack[2], a kinase conformational label has the following eight annotations:

- **Spatial Label:** Describes the position and orientation of the DFG motif as DFGin, DFGout, DFGinter, or None when key residues are not correctly mapped.
- **Dihedral Label:** Captures the backbone and side chain dihedral angles of the X-DFG motif by using the Ramachandran region annotation (A, B, L, and E) for the X, D, and F residues and the DFG-Phe $\chi_1$ rotamer (minus = $-60°$, plus = $+60°$, and trans = $180°$). These labels include BLAminus, BLBplus, ABAminus, BBAminus, BLBtrans, BLBminus, BLAplus, BABtrans, or None when key residues are not correctly mapped.
- **$\alpha$C-helix Label:** Indicates the placement of the $\alpha$C-helix as either Chelix-in, Chelix-out, or None when key residues are not correctly mapped.
- **Salt Bridge Label:** Denotes the formation of the conserved salt bridge as SaltBr-in, SaltBr-out, or None when key residues are not correctly mapped.
- **N-term. Act. Loop Label:** Classifies the N-terminal region of the activation loop as ActLoopNT-in, ActLoopNT-out, or None when key residues are not correctly mapped.
- **C-term. Act. Loop Label:** Classifies the C-terminal region of the activation loop as ActLoopCT-in, ActLoopCT-out, or None when key residues are not correctly mapped.
- **Spine Label:** Indicates the assembly state of the regulatory spine as Spine-in or Spine-out, or None when key residues are not correctly mapped.
- **Ligand Type:** Categorizes the bound ligand into Type 1, Type 1.5_Front, Type 1.5_Back, Type 2, Type 3, Allosteric, and No_ligand.

**Details of Metrics for Diversity Analysis.**

- `lys_glu_distance`: Tracks the separation between the conserved $\beta$3-Lysine and $\alpha$C-Glutamate, critical for evaluating $\alpha$C-helix packing.
- `saltbridge_distance`: Monitors the formation and structural stability of the canonical salt bridge network.
- `spine_distance`: Measures the spatial arrangement of the hydrophobic residues comprising the regulatory spine, indicating its assembly state.
- `dfg_hrd_distance`: Evaluates N-terminal activation-loop coupling by measuring the spatial separation between the DFG and catalytic HRD motifs.
- `asp_phi`: Captures the $\phi$ backbone dihedral angle of the DFG Aspartate residue.
- `asp_psi`: Captures the $\psi$ backbone dihedral angle of the DFG Aspartate residue.
- `asp_chi1`: Describes the primary side-chain torsion ($\chi_1$) of the DFG Aspartate.
- `asp_chi2`: Describes the secondary side-chain torsion ($\chi_2$) of the DFG Aspartate. This is treated with twofold ("$\pi$") symmetry so that flipped carboxylate placements are not penalized as opposite phases on the circle.
- `phe_phi`: Captures the $\phi$ backbone dihedral angle of the DFG Phenylalanine residue.
- `phe_psi`: Captures the $\psi$ backbone dihedral angle of the DFG Phenylalanine residue.
- `phe_chi1`: Describes the primary side-chain torsion ($\chi_1$) of the DFG Phenylalanine.
- `phe_chi2`: Describes the secondary side-chain torsion ($\chi_2$) of the DFG Phenylalanine. Like the Aspartate, this is treated with twofold ("$\pi$") symmetry so that flipped aromatic substituent placements are not counted as opposite phases.

### S2 Supplementary Tables

Table S1: **Percentage of KinCoRe spatial and dihedral label combinations for human kinases.** The definitions of Spatial and Dihedral Labels are defined in Section S1. From the KinCoRe server PDB inventory (https://dunbrack.fccc.edu/kincore/home) and for KinConfBench (KCB) reference chains which focuses on a subset of high quality samples and filters out None labels during selection.

| Spatial Label | Dihedral Label | All PDB | KinConfBench |
|---|---|---|---|
| DFGin | BLAminus | 6806 | 1430 |
| DFGin | BLAplus | 316 | 27 |
| DFGin | ABAminus | 993 | 183 |
| DFGin | BLBminus | 433 | 96 |
| DFGin | BLBplus | 1008 | 238 |
| DFGin | BLBtrans | 255 | 102 |
| DFGinter | BABtrans | 26 | 5 |
| DFGout | BBAminus | 677 | 144 |
| DFGin | None | 836 | 0 |
| DFGinter | None | 224 | 0 |
| DFGout | None | 365 | 0 |
| None | None | 413 | 0 |

Table S2: **Distribution of KinCoRe spatial and dihedral labels for KinConfBench (KCB) and cofolding model predictions.** KCB percentages are derived from the 2,225-chain set described in the Results. Percentages for Boltz-2 ($n = 28,400$), Chai-1 ($n = 28,399$), and Protenix ($n = 28,400$) are calculated across 1,420 distinct kinase systems. * The small percentage of "None" labels results from residue mismatches during the KinCoRe labeling process.

| Spatial | Dihedral | KCB (%) | Boltz-2 (%) | Chai-1 (%) | Protenix (%) |
|---|---|---|---|---|---|
| DFGin | BLAminus | 64.3 | 65.8 | 77.5 | 65.1 |
| DFGin | BLAplus | 1.2 | 1.1 | 1.2 | 1.5 |
| DFGin | ABAminus | 8.2 | 3.9 | 4.8 | 3.3 |
| DFGin | BLBminus | 4.3 | 3.0 | 2.0 | 3.4 |
| DFGin | BLBplus | 10.7 | 8.0 | 8.5 | 8.3 |
| DFGin | BLBtrans | 4.6 | 11.6 | 0.1 | 11.7 |
| DFGinter | BABtrans | 0.2 | 0.0 | 0.0 | 0.0 |
| DFGout | BBAminus | 6.5 | 5.3 | 4.3 | 5.4 |
| DFGin | None* | 0.0 | 0.4 | 0.4 | 0.2 |
| DFGinter | None* | 0.0 | 0.8 | 0.6 | 0.1 |
| DFGout | None* | 0.0 | 0.1 | 0.5 | 0.1 |
| None* | None* | 0.0 | 0.0 | 0.1 | 1.0 |

Table S3: **Conformational (KinCoRe label) summary on 950 systems passing geometric filters.** Systems ≥1 correct: systems with at least one all-labels-correct prediction in top-20. Avg correct/system: mean KinCoRe-correct predictions per system (out of top-20).

| Model | Systems ≥1 correct | Avg correct/system |
|---|---|---|
| Boltz-2 | 779 (82.0%) | 14.2 |
| Chai-1 | 724 (76.2%) | 12.4 |
| Protenix | 776 (81.7%) | 14.1 |

Table S4: **110 systems where all cofolding predictions fail to generate the correct conformation as the ground truth.** Each entry is a KinConfBench system key (`GENE_Name_Ligands`).

| **System identifiers** | | | | |
|---|---|---|---|---|
| ABL1_HUMAN_66K | ABL1_HUMAN_STI_1N1 | ACK1_HUMAN_1G0 | ACK1_HUMAN_DBQ | ACK1_HUMAN_LWX |
| ACK1_HUMAN_R7P | ACK1_HUMAN_T74 | ACK1_HUMAN_T95 | ACK1_HUMAN_WTP | ALK_HUMAN_25J |
| ALK_HUMAN_6YL | ALK_HUMAN_AWJ | ALK_HUMAN_HKJ | ALK_HUMAN_J3Y | AURKA_HUMAN_9YQ |
| AURKA_HUMAN_X6D | BRAF_HUMAN_5I4 | BRAF_HUMAN_FP3 | BRAF_HUMAN_P06 | BTK_HUMAN_YDA |
| CDK1_HUMAN_1QK | CDK2_HUMAN_02Z | CDK2_HUMAN_06Z | CDK2_HUMAN_09Z | CDK2_HUMAN_0S0 |
| CDK2_HUMAN_106 | CDK2_HUMAN_20K | CDK2_HUMAN_26Z | CDK2_HUMAN_A1A1H | CDK2_HUMAN_A1D6S |
| CDK2_HUMAN_A27 | CDK2_HUMAN_DTQ | CDK2_HUMAN_ES4 | CDK2_HUMAN_JWS | CDK2_HUMAN_LS1 |
| CDK2_HUMAN_LS3 | CDK2_HUMAN_LS4 | CDK2_HUMAN_R0N | CDK2_HUMAN_SU9 | CDK2_HUMAN_WQ6 |
| CDK2_HUMAN_X02 | CDK2_HUMAN_X06 | CDK2_HUMAN_X19 | CDK2_HUMAN_X35 | CDK2_HUMAN_X36 |
| CDK2_HUMAN_X3A | CDK2_HUMAN_X40 | CDK2_HUMAN_X42 | CDK2_HUMAN_X43 | CDK2_HUMAN_X44 |
| CDK2_HUMAN_X62 | CDK2_HUMAN_X6B | CDK2_HUMAN_Y8L | CDK2_HUMAN_Z19 | CDK2_HUMAN_Z63 |
| CDK2_HUMAN_Z71 | CDK5_HUMAN_65L | CDK6_HUMAN_24V | CDK6_HUMAN_AP9 | CDK6_HUMAN_LQQ |
| CDK7_HUMAN_I73 | CDK7_HUMAN_WZ8 | CDK8_HUMAN_C1I | CHK1_HUMAN_306 | CHK1_HUMAN_373 |
| CHK1_HUMAN_76A | CHK1_HUMAN_H0K | CHK1_HUMAN_YM6 | CLK1_HUMAN_Q7K | CLK1_HUMAN_WAZ |
| DAPK1_HUMAN_BD4 | DAPK1_HUMAN_LU2 | DAPK1_HUMAN_PIT | DAPK1_HUMAN_STU | DAPK3_HUMAN_4RB |
| DCLK1_HUMAN_XBD | EPHA2_HUMAN_DXK | EPHA2_HUMAN_L66 | EPHA2_HUMAN_QRD | EPHA2_HUMAN_QRR |
| EPHA2_HUMAN_WT3 | IGF1R_HUMAN_PDR | JAK3_HUMAN_79T | KC1D_HUMAN_AUE | KCC2D_HUMAN_K88 |
| KSYK_HUMAN_1B6 | KSYK_HUMAN_4MG | KSYK_HUMAN_685 | KSYK_HUMAN_X7G | M3K5_HUMAN_NJV |
| M3K5_HUMAN_STU | M4K1_HUMAN_2WI | M4K1_HUMAN_A1AP0 | M4K1_HUMAN_YK1 | MK01_HUMAN_2SH |
| MK01_HUMAN_33A | MK01_HUMAN_35X | MK01_HUMAN_362 | MK01_HUMAN_5ID | MK01_HUMAN_F29 |
| MK01_HUMAN_FRZ | MK07_HUMAN_R4L | MK14_HUMAN_GK1 | SRPK1_HUMAN_RXZ | VRK2_HUMAN_7DZ |
| VRK2_HUMAN_KJD | WEE1_HUMAN_34W | WEE1_HUMAN_99J | WEE1_HUMAN_99M | WEE1_HUMAN_99V |

#### S3 Supplementary Figures

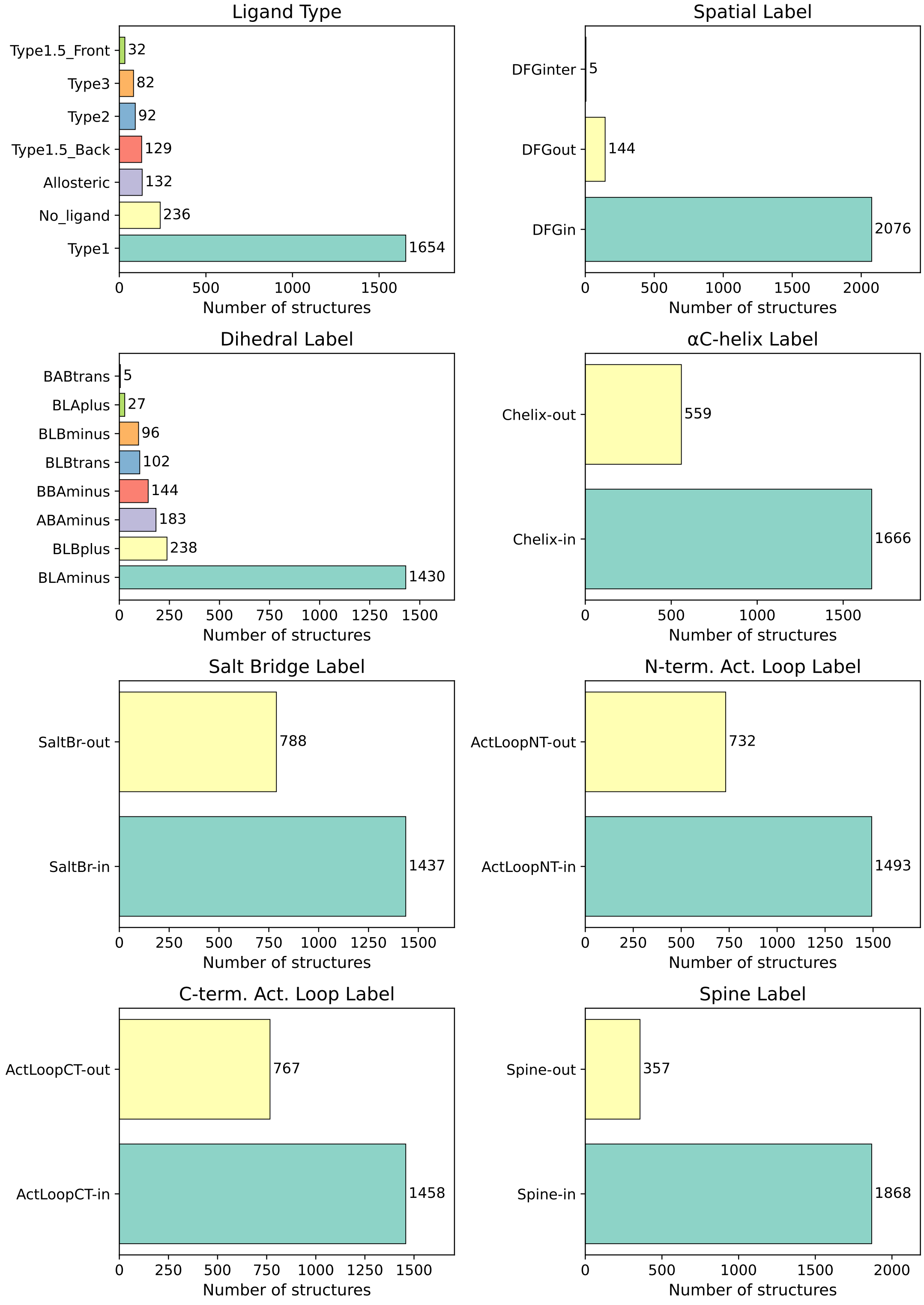


Figure S1: **KinConfBench curation and composition.** Summary of how the benchmark is built from PDB-derived kinase chains: counts by holo versus apo, single- versus multi-ligand complexes, redundancy filtering, and the distribution of entries across Manning kinase groups and ligand/chemotype categories. Together these panels document that the final benchmark is holo-heavy (reflecting the PDB) yet retains diverse genes and chemotypes, and that reported model statistics are computed on the curated non-redundant set rather than the raw PDB pull.

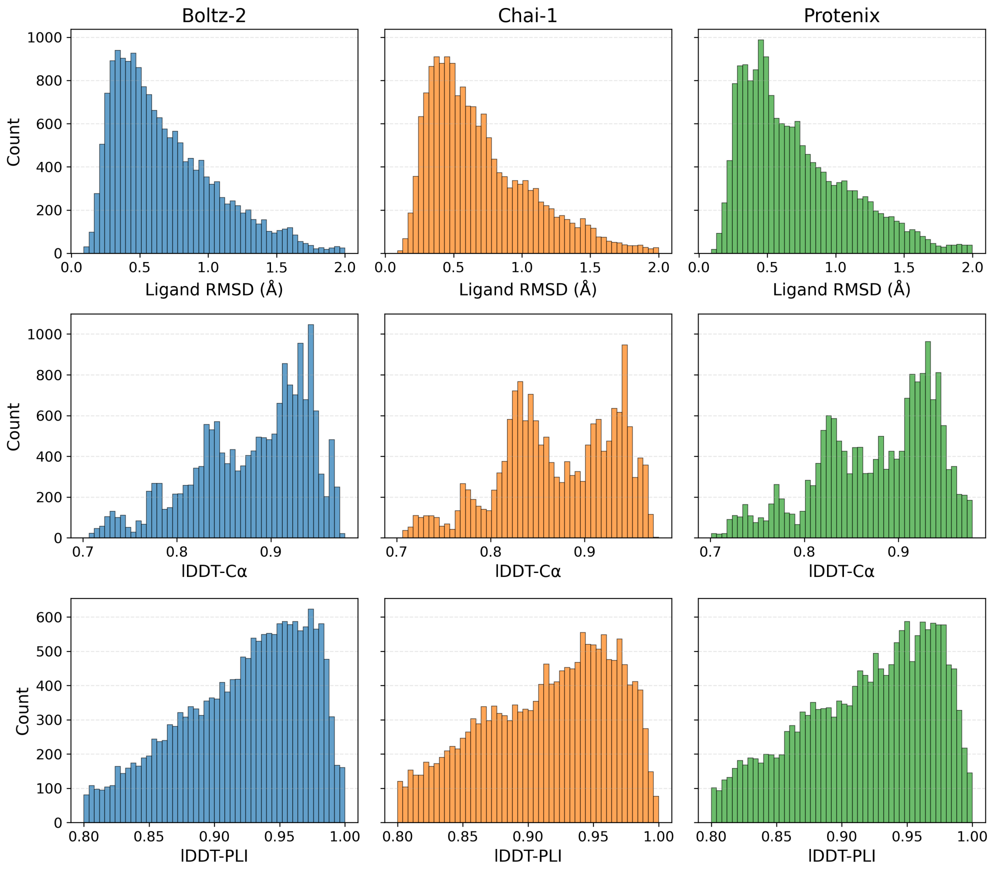


Figure S2: **Distribution of geometric metrics for cofolding predictions.** Distributions of ligand heavy-atom RMSD, lDDT-C$\alpha$, and lDDT-PLI for ensemble predictions that pass the benchmark geometric filters (lDDT-C$\alpha \geq 0.7$, lDDT-PLI $\geq 0.8$, ligand RMSD $< 2$ Å). We have 950 protein-ligand kinase systems from Boltz-2 (Blue, 17,409 samples), Chai-1 (Orange, 16,471 samples), and Protenix (Green, 17,125 samples).

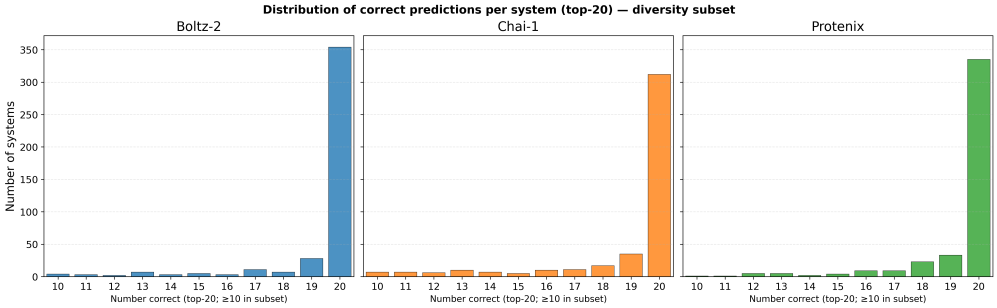


Figure S3: **Correct-prediction counts on the high-yield diversity subset.** Same quantity as in Figure 2(c), but restricted to 509 systems where Boltz-2, Chai-1, and Protenix each have at least ten all-labels-correct predictions in the top-20, the subset used for ensemble diversity analysis. The horizontal range is truncated to 10-20 so that differences in the upper tail (near-enumeration of the correct state) are visible. A single-row layout with a shared $y$-axis facilitates direct comparison of how densely each model covers the correct label manifold when it is readily accessible.

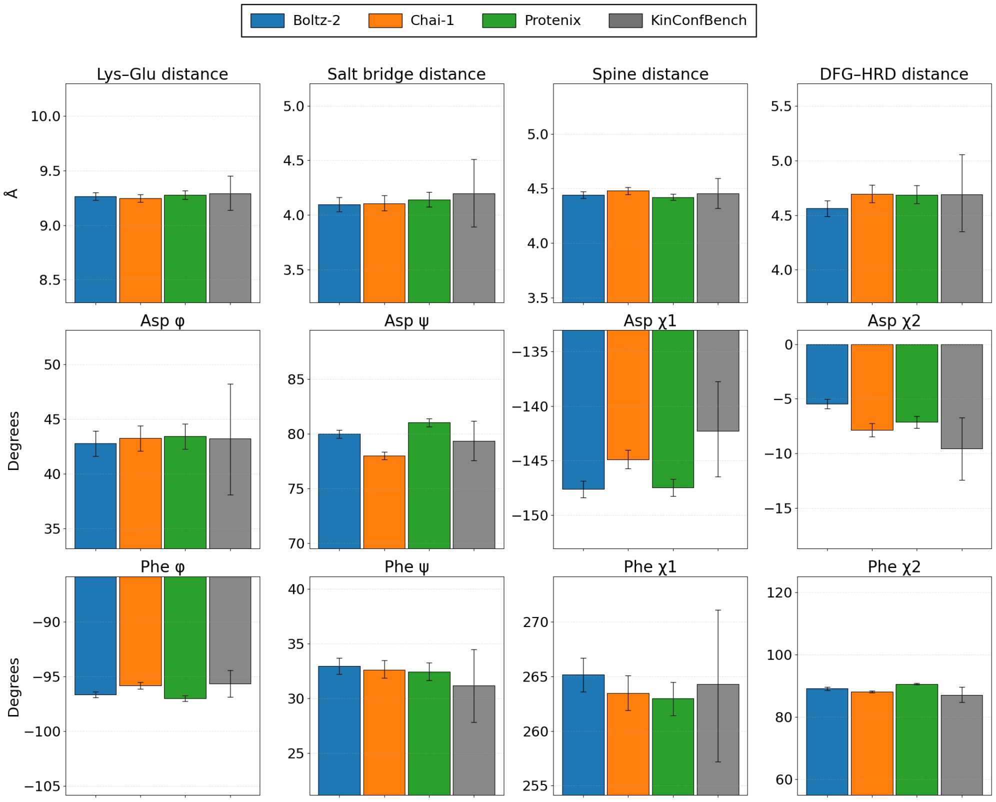


Figure S4: **Mean of key distance and angle metrics for cofolding predictions and KinConfBench.** The plot displays mean values for Boltz-2 (blue), Chai-1 (orange), and Protenix (green) predictions, with KinConfBench PDB reference values provided in gray for comparison. Error bars are bootstrap 95% confidence intervals for each pooled mean.

### References

[1] Manning G, Whyte DB, Martinez R, Hunter T, Sudarsanam S. The Protein Kinase Complement of the Human Genome. Science. 2002 Dec;298(5600):1912-34. Available from: https://www.science.org/doi/10.1126/science.1075762.

[2] Modi V, Dunbrack RL. Kincore: a web resource for structural classification of protein kinases and their inhibitors. Nucleic Acids Research. 2022 Jan;50(D1):D654-64. Available from: https://academic.oup.com/nar/article/50/D1/D654/6395339.
